# Molecular docking: Bioactive compounds of *Mimosa pudica* as an inhibitor of *Candida albicans* Sap 3

**DOI:** 10.1101/2022.09.06.506736

**Authors:** Gusnia Meilin Gholam, Iman Akhyar Firdausy, I. Made Artika, Ramadhani Malik Abdillah, Ridwan Putra Firmansyah

**Author notes:** Corresponding author (G.M.G).

## Abstract

*Candida albicans* (*C. albicans*) is a commensal microbiota that resides in humans. However, in certain cases, C. albicans can infect and cause several diseases to humans. This study aimed to investigate the interaction between Mimosa pudica bioactive compounds and *C. albicans* Sap 3. Molecular docking analysis was carried out using YASARA structure. The procedures involved preparation of ligands and target receptor, molecular docking, data analysis and visualization. All 3D ligands were downloaded from PubChem NCBI, while target receptor was downloaded from RCSB PDB. The interaction between Mimosa pudica bioactive compounds against Sap 3 resulted in a binding energies ranges from 5,168 – 7,480 kcal/mol and most of the interactions formed were relatively strong. Furthermore, the test ligands had contact with the catalytic residues and substrate binding site pockets S1/S2/S3/S4 on the target receptor. Bioactive compounds of Mimosa pudica have relatively good interactions in inhibiting *C. albicans* Sap 3.

## 1. Introduction

Most of the human pathogenic fungi of the genus *Candida* reside in animals and humans ^1^. Invasive *Candida* infection is one of the most common fungal infections globally ^2,3^. In western countries, individuals generally have *Candida* as a commensal microbiota in their gut ^1^. In the United States, *Candida* spp. is reported to be one of the leading causes of healthcare-associated infections ^2,3^. *Candida* infection is also often associated with medical devices such as central venous catheters, cardiovascular machines, and urinary catheters ^2,4^. Among the species of *Candida* sp., *Candida albicans* (*C. albicans*) (37%) is the most commonly found in the clinical species ^2^. *C. albicans* is known as a commensal microbiota that lives in normal humans. However, *C. albicans* is also one of the most common fungal species causing opportunistic infections, such as candidiasis ranging from superficially invasive infections to life-threatening in debilitated patients ^5,6^. Invasive candidiasis is associated with a high mortality rate, ranging from 20 to 49%. *C. albicans* can be found to colonize various mucosal surfaces such as skin, mouth, and vagina; most studies consider the gastrointestinal tract as the main entry point for *C. albicans* to enter the bloodstream ^5–9^. On the other hand, *C. albicans* is the most common causative agent of oral, vaginal, and disseminated candidiasis. Oral candidiasis is one of the most common opportunistic infectious diseases among patients suffering from HIV infection. *C. albicans* can multiply rapidly, invade tissues, and cause symptomatic mucosal lesions in people with immune systems disorder ^10–12^. Therefore, good immune systems are important to maintain the fungus in a commensal state, preventing invasion, epithelial damage, and mucosal infection ^1,13^. It is estimated that more than 7.5 million people globally have been infected by invasive candidiasis. There are unwanted side effects, ineffectiveness, and the rapid development of resistance by fungi, therefore there is a need for the development of new antifungals ^9^. Many factors and activities have been identified as the cause of the pathogenicity of *C. albicans* ^1,14^, secreted aspartic proteinases (Sap) 1-3 have been observed to play an important role in adhesion and tissue damage in local infections ^15,16^. In-depth understanding of the contribution of the Sap family to the pathogenicity of *C. albicans*, obtained by apprehending the *C. albicans* strains; Sap 1-3 are involved in mucosal infections, and Sap 4-6 are involved in systemic infections ^15^. *C. albicans* Sap plays a multimodal role in the infection process, therefore the development of inhibiting Sap as targets for a wide variety of infections caused by *C. albicans* appears to be a good strategy ^10^.

*Mimosa pudica* Linn (*M. pudica*) is a medicinal plant with pharmaceutical and nutraceutical potential; this plant is also popular among traditional healers to treat various diseases ^17^. Herbal plants have been widely used to treat several diseases and are used as traditional medicine ^18^. This plant was observed because of its thigmotactic and seismonastic movements. *M. pudica* is known for its analgesic, anti-inflammatory, diuretic activity, insomnia, and urogenital infections ^17,19–21^. Phytochemical compounds in *M. pudica* are shown in the study of Vijayalakshmi et al. ^17^.

The molecular docking method is one of the *in silico* research methods and is one of the most powerful techniques to help discover new ligands for proteins with known structures, thus playing a pivotal role in structure-based drug design ^22^. Therefore, through this study, we investigated the possibility of an interaction between bioactive compounds from *M. pudica* and the Sap 3 *Candida albicans* to find new inhibitor candidates.

## 2. Materials and Methods

### 2.1 Receptor preparation

Sap 3 with the code “2H6T” was downloaded from the RCSB PDB website (https://www.rcsb.org/, accessed on 13 July 2022). Preparation was carried out using YASARA structure (Bioinformatics 30.2981-2982 Version 19.9.17) by adding hydrogen atoms, removing water molecules around the receptor, and adjusting the receptor pH value to 7.40 ^23^.

### 2.2 Ligand preparation

The bioactive compounds of *M. pudica* was obtained from Vijayalakshmi et al. ^17^, then the 3D structure was downloaded from the NCBI PubChem page (https://pubchem.ncbi.nlm.nih.gov/, accessed on 13 July 2022) ^24^. The preparation also uses YASARA structure to add hydrogen atoms, adjust the pH value to 7.40, and energy minimization ^23,25^.

### 2.3 Molecular docking

To explore the possibility of ligand binding into the pocket area of the substrate binding site or receptor catalytic residue, molecular docking was carried out using YASARA structure software ^26^. YASARA structure was set to AutoDock Vina settings, AMBER03 force field, and 25 runs (dock_run). The docking area was set to around all atoms to expand the docking area. Then the other parameters remained unchanged. Finally, the resulting conformers were analyzed to determine hydrogen bonding, hydrophobic interactions, electrostatic interactions, and other interactions ^23,27^.

### 2.4 Data analysis and visualization

After docking, the receptor-ligand complex interactions were analyzed using BIOVIA Discovery Studio Visualizer v21.1.0.20298. The results of the analysis were in the form of 2D and 3D structures to help visualize and analyze the interaction pattern of the ligand-protein complex ^9^.

## 3. Results and Discussion

This research used the molecular docking method to bind compounds (test ligands) from *Mimosa pudica* (*M. pudica*) to Sap 3 *Candida albicans* (*C. albicans*). The docking process did not target specific residues or limit the docking area, therefore it is hoped that when the tethering process took place, the ligand can have more comprehensive residual contact and still have contact with essential residues of the target receptor.

It is widely known that *C. albicans* has Sap 1 to Sap 10 genes. Sap is one of the classic virulent factors whose expression is modulated by several conditions such as the influence of pH, temperature, site of infection, and physicochemical environmental conditions ^10,28^. This research used one of the Saps, namely Sap 3. We chose Sap 3 with PDB ID code 2H6T because it has been crystallized with pepstatin ^15^. The pepstatin used as the inhibitor standard to refer to the standard binding energy, dissociation constant (Kd), and amino acid residues ^10^.

Based on the results of this study, interestingly, pepstatin had the highest binding energy of 8.857 kcal/mol and Kd 321908.8438 pM. The test ligands that have binding energy close to pepstatin are turgorin with a binding energy of 7,480 kcal/mol and *K*_d_ 3289177.500 pM, then based on the binding energy respectively, D-glucuronic acid with a binding energy of 6.219 kcal/mol and Kd 27632016.000 pM, gallic acid with a binding energy of 6.053 kcal/mol and Kd 36567240.000 pM, L-norepinephrine with a binding energy of 5.770 kcal/mol and Kd 58956804.000 pM, mimosine with a binding energy of 5.599 kcal/mol and Kd 78682576.000 pM, L-ascorbic acid with a binding energy of 5.390 kcal/mol and Kd 111963792.000 pM, and the lowest binding energy is linolenic acid with a binding energy of 5.168 kcal/mol and Kd 162856752.000. The more positive value of the binding energy indicates stronger bond between the ligand and the target receptor whereas ^29,30^. The results of the binding energies are presented in Figure 1. The smaller value of Kd indicates that ligand binding affinity againts the receptor is stronger ^31–33^. We also show the Kd and overall residual contacts in Table 1.

**Figure 1.**
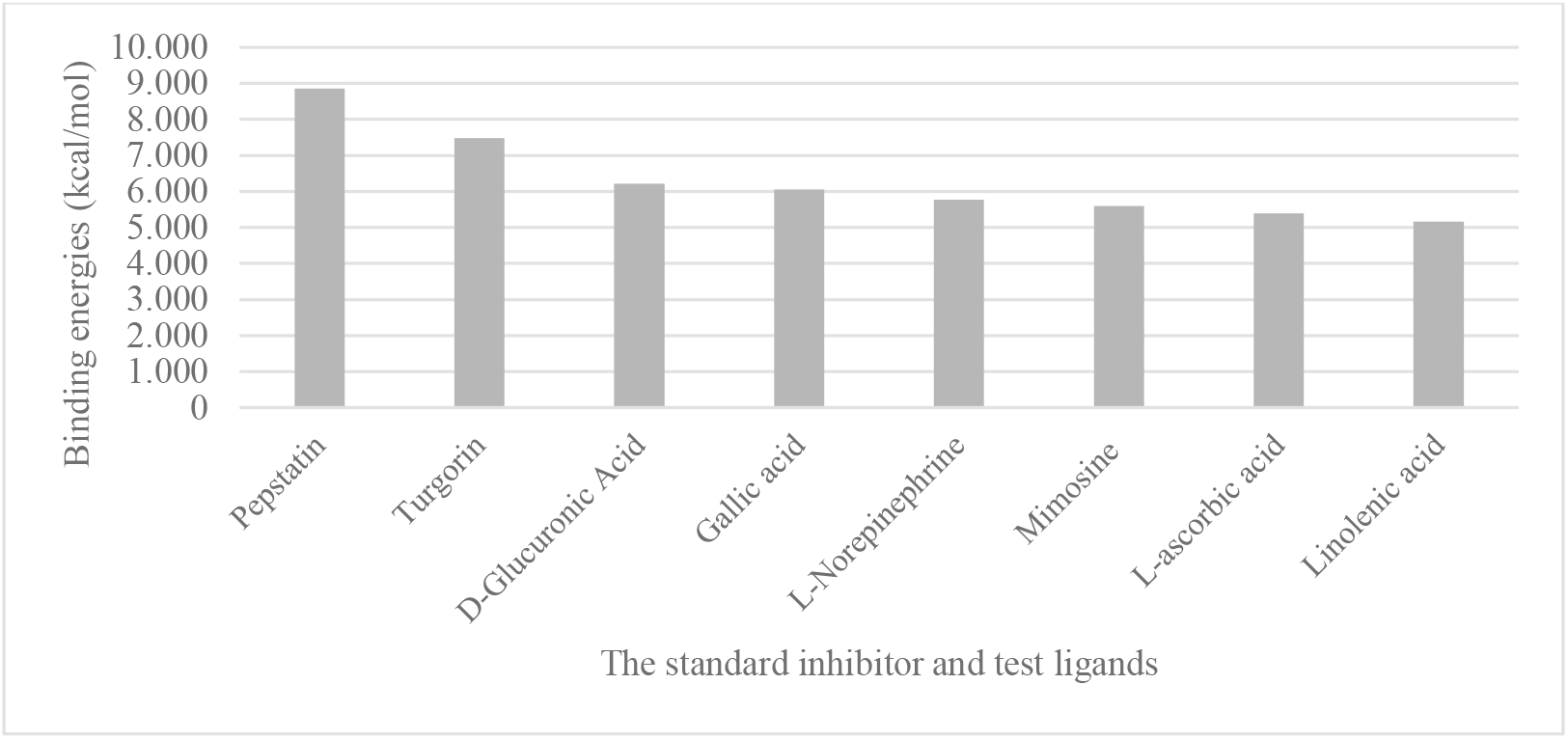
The binding energies of *M. pudica* bioactive compounds against Sap 3 determined using YASARA structure

**Tabel 1.** Molecular docking results of *M. pudica* bioactive compounds against Sap 3 using YASARA structure

| Ligand | Dissoc. Constant (pM) | Contacting receptor residues |
| --- | --- | --- |
| Pepstatin (standard inhibitor) | 321908.8438 | A VAL 12 A SER 13 A VAL 30 A ASP 32 A THR 33 A GLY 34 A SER 35 A SER 36 A ILE 82 A GLU 83 A TYR 84 A GLY 85 A ASP 86 A VAL 119 A ASP 120 A GLN 121 A ILE 123 A GLY 127 A TYR 128 A ASN 192 A GLU 193 A LEU 194 A ARG 195 A LEU 216 A ASP 218 A GLY 220 A THR 221 A THR 222 A ILE 223 A TYR 225 A TYR 303 A ILE 305 |
| Turgorin | 3289177.500 | A VAL 30 A ASP 32 A GLY 34 A GLU 83 A TYR 84 A GLY 85 A ASP 86 A THR 88 A VAL 119 A ILE 123 A ARG 195 A LEU 216 A ASP 218 A GLY 220 A THR 221 A TYR 303 A ILE 305 |
| D-Glucuronic Acid | 27632016.000 | A ASN 9 A GLN 11 A VAL 12 A TYR 14 A ARG 162 A GLN 163 A THR 222 A ALA 281 A LEU 297 A ASN 309 A ARG 312 |
| Gallic acid | 36567240.000 | A ASN 9 A GLN 11 A TYR 14 A ARG 162 A GLN 163 A THR 222 A LEU 297 A ASN 309 A ARG 312 |
| L-Norepinephrine | 58956804.000 | A VAL 30 A ASP 32 A GLY 34 A TYR 84 A GLY 85 A ASP 86 A THR 88 A VAL 119 A ILE 123 A ASP 218 A GLY 220 A THR 221 A ILE 305 |
| Mimosine | 78682576.000 | A SER 13 A VAL 30 A ASP 32 A GLY 34 A SER 35 A TYR 84 A GLY 85 A ASP 86 A THR 88 A VAL 119 A ILE 123 A ASP 218 A GLY 220 A THR 221 |
| L-ascorbic acid | 111963792.000 | A THR 33 A GLY 34 A SER 35 A SER 36 A GLY 127 A TYR 128 A LYS 129 A ASN 192 A GLU 193 A LEU 194 A LEU 216 |
| Linolenic acid | 162856752.000 | A ASP 32 A GLY 34 A SER 35 A TYR 84 A GLY 85 A ASP 86 A LEU 216 A ASP 218 A GLY 220 A THR 221 A THR 222 A ILE 223 A TYR 225 A ILE 305 |

It is important to note that through Borelli et al.^15^ research, Sap 3 was known to be divided into several pockets of substrate binding sites on S1, S2, S3, and S4. S1 consists of VAL30, TYR84, ASP86, THR88, VAL119, and ILE123 amino acid residues. The S2 consists of GLY85, ASP86, THR221, TYR225, SER301, TYR303, and ILE305 amino acid residues. S3 consists of VAL12, SER13, ASP86, THR88, SER188, ASP120, and GLY220 amino acid residues. S4 consists of VAL12, THR222, ILE223, TYR225, GLN295, LEU297, and GLY299 amino acid residues, while the catalytic residues are located in ASP32 and ASP218. This study usedthese important residues as a criterion.

Through the help of the Discovery Studio software, the patterns of interactions that are formed within the complex can be seen. The results of the analysis are presented in Table 2.

**Tabel 2.**
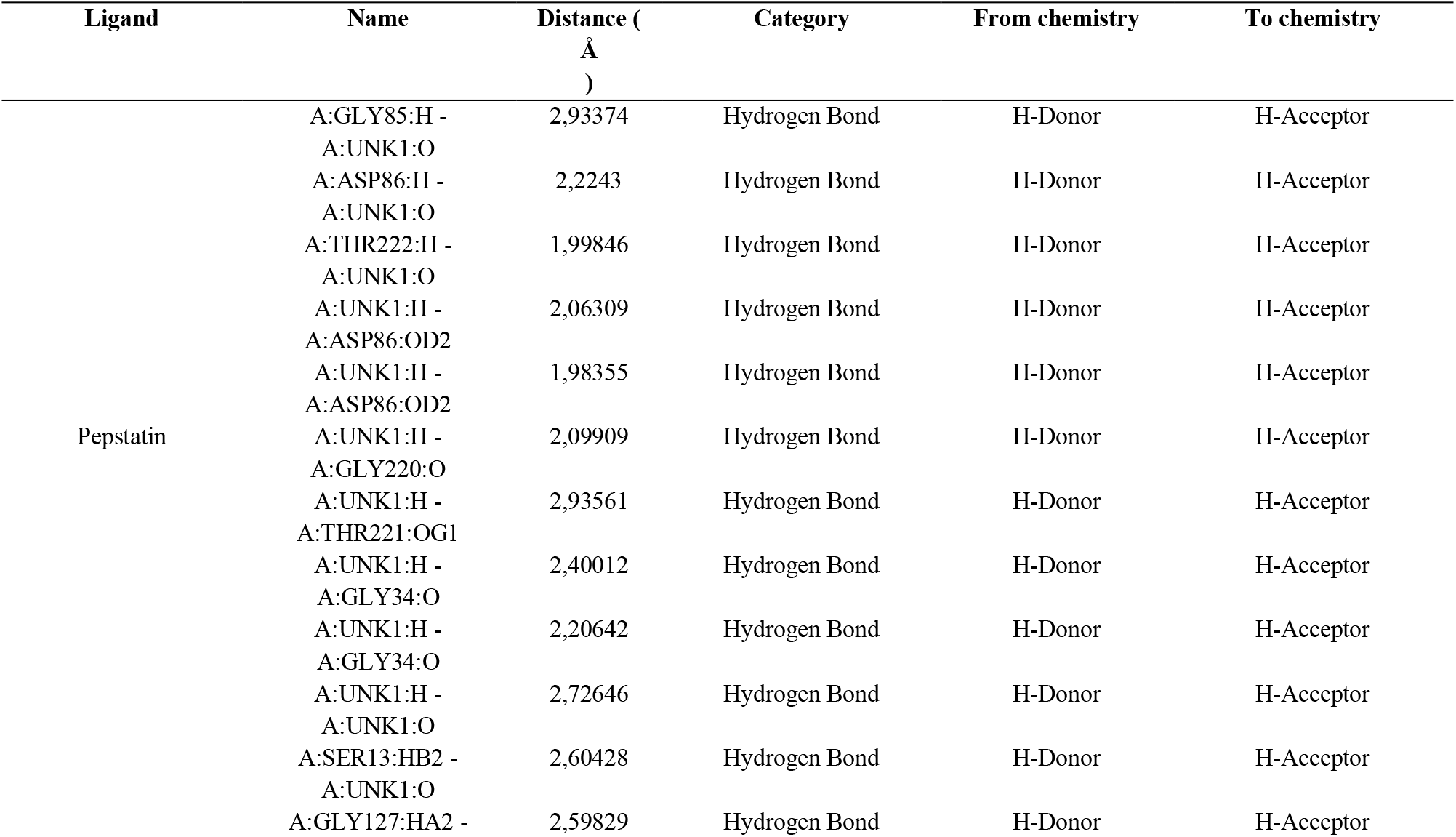
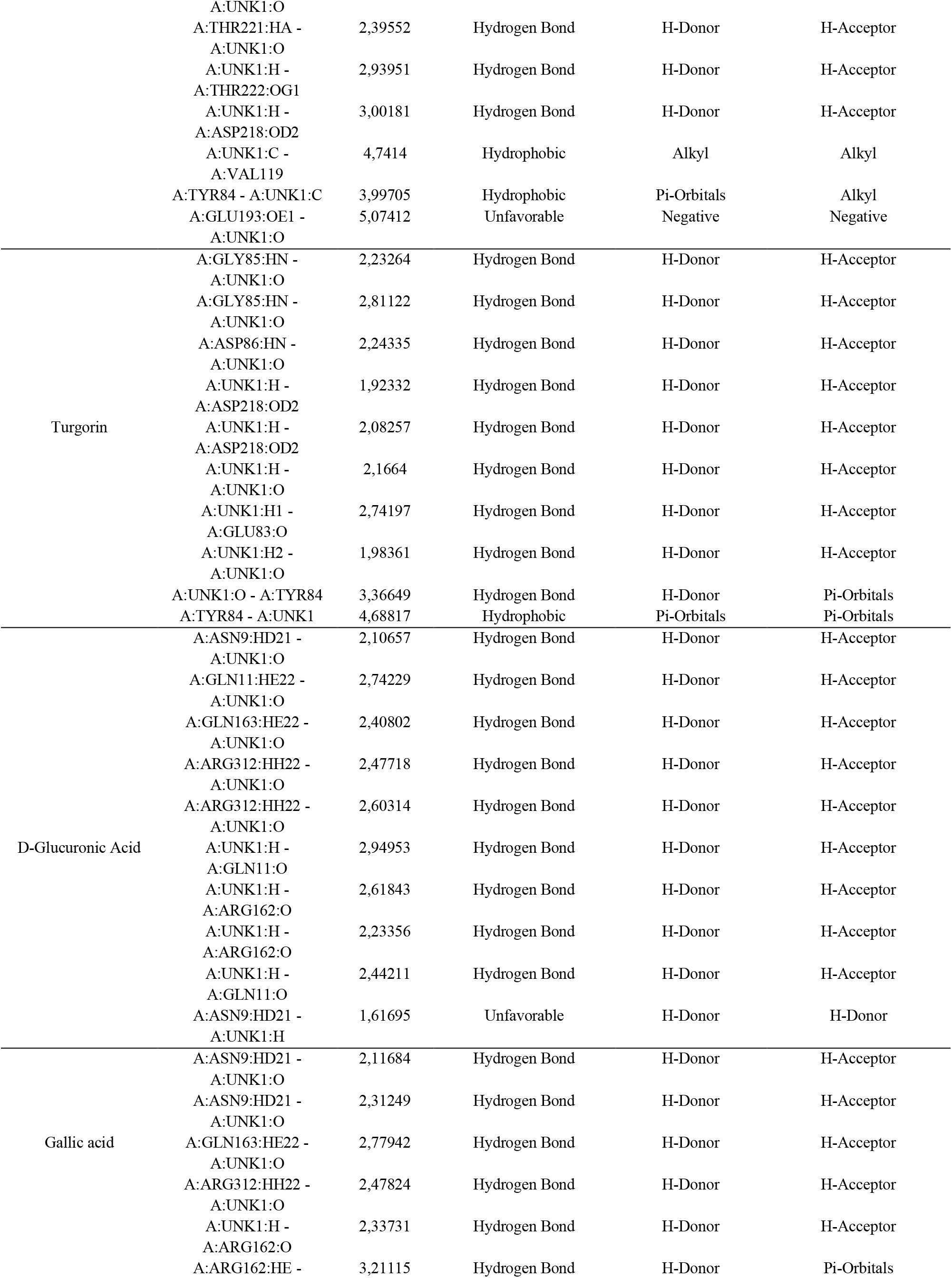
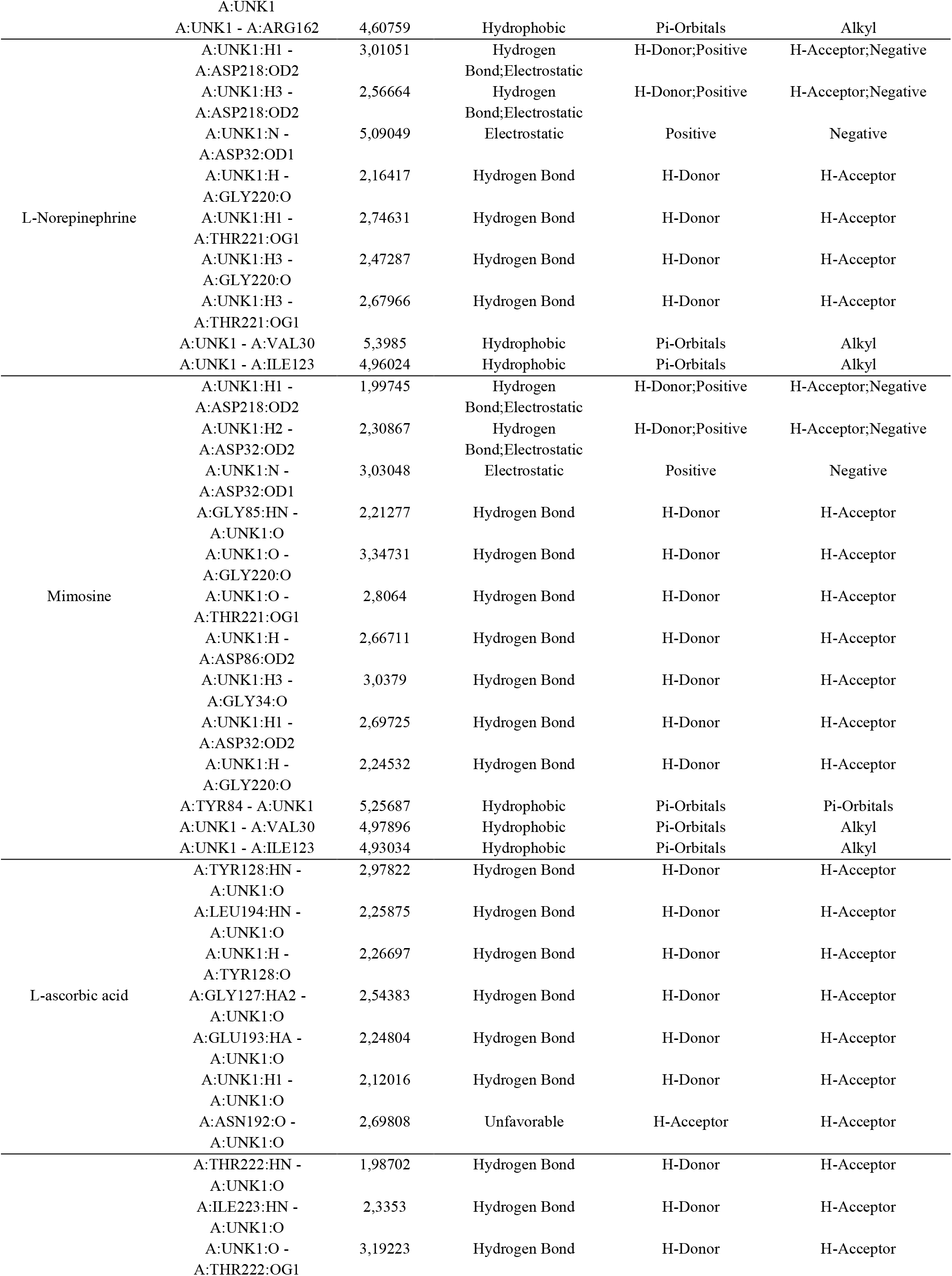
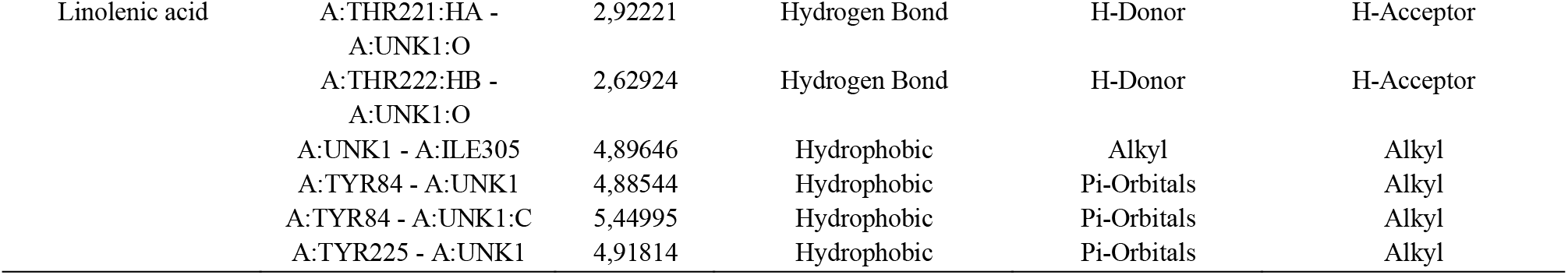
The interactions of *M. pudica* bioactive compounds against *C. albicans* Sap 3

Pepstatin tends to make hydrogen bonds with the target receptor (Figure 2). Hydrogen bonds have a distance ranging between 1.99846 - 3.00181 Å. The category of hydrogen bonds formed in the Sap3-Pepstain complex has from chemistry H-donor and to chemistry H-acceptor. SER13, GLY85, ASP86, and THR222 bind with the O atom in pepstatin. GLY220 and THR222 bind with the H atom of pepstatin. Pepstatin also has hydrophobic and unfavorable interactions. The hydrophobic interaction between A:TYR84 - A:UNK1:C was Pi-Alkyl type. Other hydrophobic interactions also formed in Alkyl type between A:UNK1:C - A:VAL119. This study found an unfavorable interaction in the Sap 3-Pepstatin complex at A:GLU193:OE1 - A:UNK1:O under the negative-negative unfavorable type. Pepstatin interacts with amino acid residues at S1/S2/S3/S4 and catalytic residues. In Borelli et al.^15^ study, pepstatin interacted with amino acid residues located at S1/S2/S3/S4. However, pepstatin suboptimally filled several binding sites, mainly at S3/S4 ^15^ and his research also explained that pepstatin is a pentapeptide produced by the *Streptomyces* ^15^.

**Figure 2.**
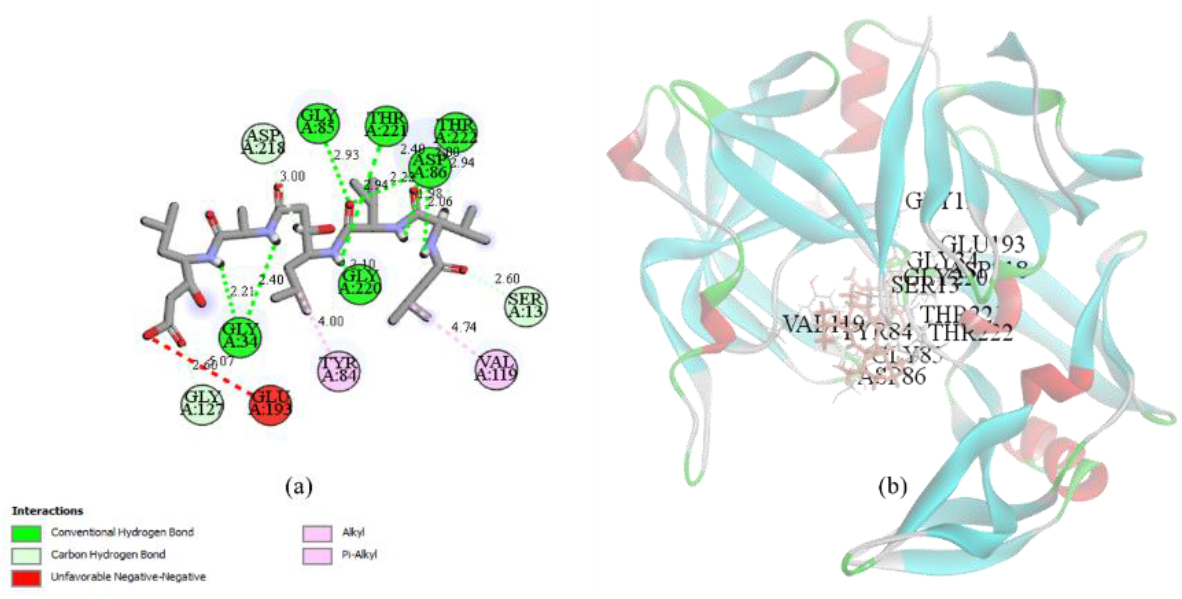
Visualization using BIOVIA Discovery Studio shows the interaction formed on the pepstatin complex with Sap 3 (a) 2D visualization shows bond distance, amino acid residues, and interaction type (b) 3D visualization shows the semi-transparent complex

Turgorin formed ten interactions with the target receptor; out of ten interactions, only one was hydrophobic (Figure 3). The hydrogen bonds formed have a distance between 1.92332 - 2.81122 Å. This ligand had contact with the ASP218 catalytic residue formed the OH group. Amino acid residues that interact with the ligand O atom were ASP85 and ASP86 by forming hydrogen bonds. TYR84 interaced with the ligand aromatic ring with a distance of 4.68817 Å, with from chemistry Pi-Orbitals and to chemistry Pi-Orbitals. The strongest hydrogen bond was found on the ASP218 catalytic residue and the ligand OH group with a distance of 1.92332 Å. These ligands interacedt with amino acid residues in S1/S2 and catalytic residues.

**Figure 3.**
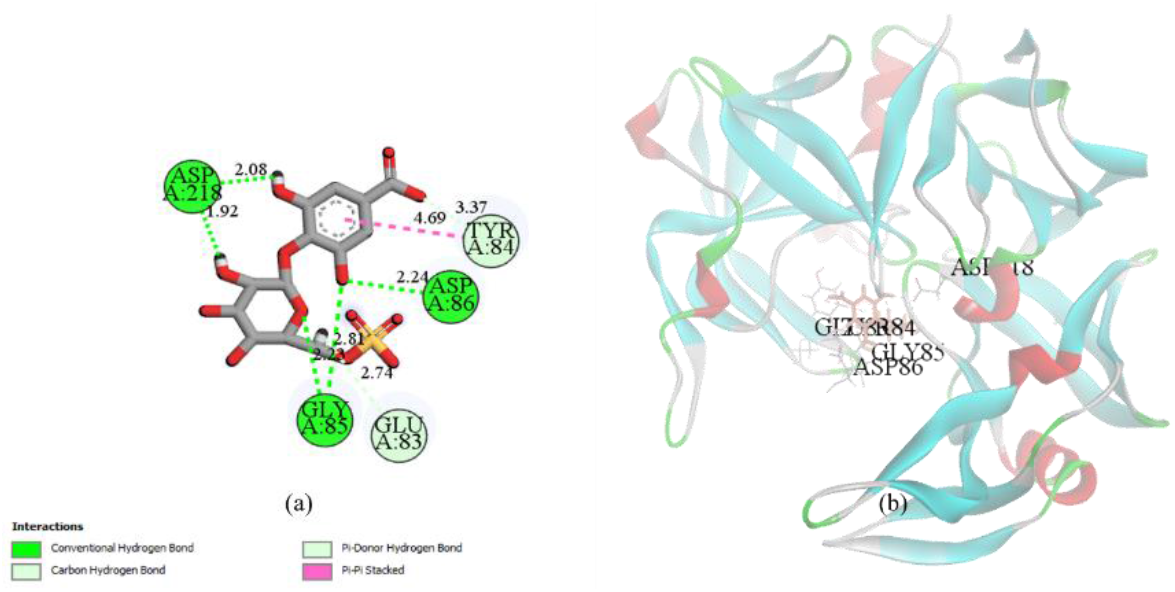
Visualization using BIOVIA Discovery Studio shows the interaction formed in the turgorine-Sap 3 complex (a) 2D visualization shows bond distance, amino acid residues, and interaction type (b) 3D visualization shows the complex in a semi-transparent manner

D-glucuronic acid, gallic acid, and L-ascorbic acid had no interaction with the substrate binding site pocket. They did not even interact with the Sap 3 target receptor’s catalytic residue (Table 2).

D-glucuronic acid had ten interactions, with details of nine hydrogen bonds and one unfavorable interaction (Figure 4). Amino acid residues that formed OH groups with ligands are ASN9, GLN11, ARG162, GLN163, and ARG312. The distance created due to the presence of hydrogen bonds ranged from 2.10657 - 2.94953 Å. Unfavorable bonds were formed on the ASN9 amino acid residues with ligand H atoms, from chemistry H-donor and to chemistry H-donor, and the distance is 1.61695 Å.

**Figure 4.**
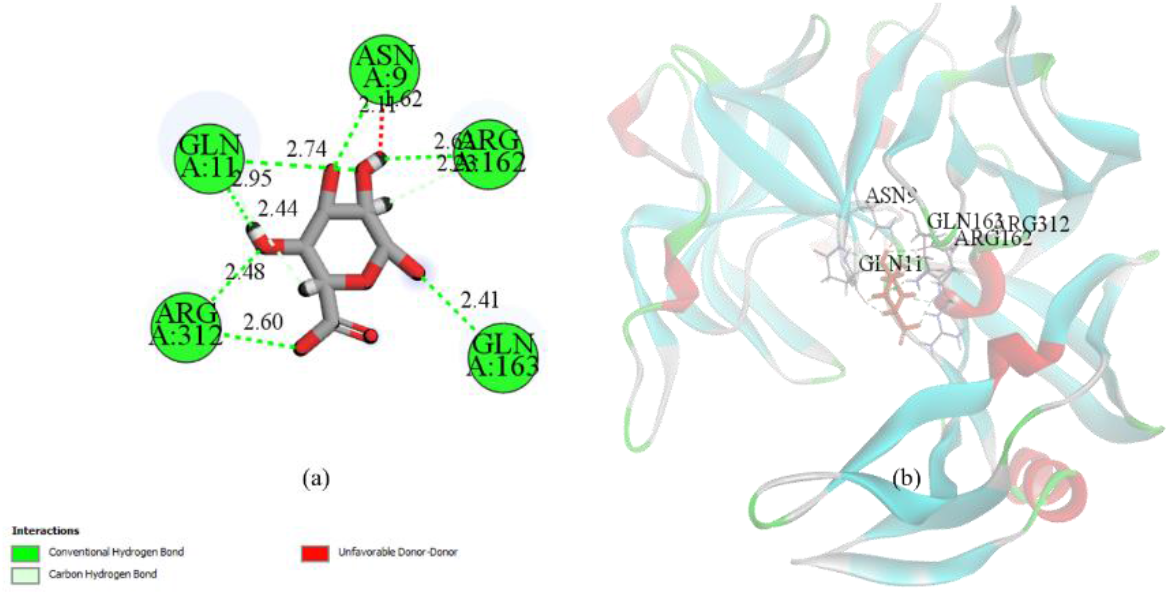
Visualization using BIOVIA Discovery Studio shows the interaction formed on the D-glucuronic acid-Sap 3 complex (a) 2D visualization shows bond distance, amino acid residues, and interaction type (b) 3D visualization shows the semi-transparent complex

Abu-Izneid et al. ^34^ explained that glucuronide or glucuronoside, which are D-glucuronic acid and glucuronide derivatives, is an important class of active pharmaceutical compounds known to have antiviral activity against viruses, such as A/H1N1 and A/H3N2 strains of influenza virus.

Through docking simulations, we looked for whether there was a possibility that D-glucuronic acid ligand has antifungal activity. From the results of the analysis of D-glucuronic acid ligand targeting Sap 3, which showed that nine hydrogen bonds were formed in the receptor-ligand complex, although this ligand had no contact with the catalytic residue or the pocket of the substrate binding site, we predicted this ligand was quite good at inhibiting the *C. albicans* Sap 3.

ARG162 formed a hydrogen bond by interacting with the aromatic ring of the gallic acid ligand with a distance of 3.21115 Å (Figure 5), and from chemistry H-donor, to chemistry Pi-orbitals. The distance produced by the presence of hydrogen bonds ranges from 2.11684 - 3.21115 Å, while the hydrophobic interaction has a distance of 4.60759 Å. Liberato et al. ^35^ through an in vitro study, explained that gallic acid has potential as antifungal activity against *Candida* spp. Another study also reported, through *in vitro* and *in silico* studies by Uma Maheshwari Nallal et al. ^30^, that gallic acid, which is classified as an active phytochemical compound, is predicted to be an effective inhibitory agent of Sap from *Candida* species. Therefore, even though in this research gallic acid did not interact with the substrate binding site pocket or the catalytic residue of the target receptor, gallic acid was predicted to have a reasonably good interaction in inhibiting Sap 3.

**Figure 5.**
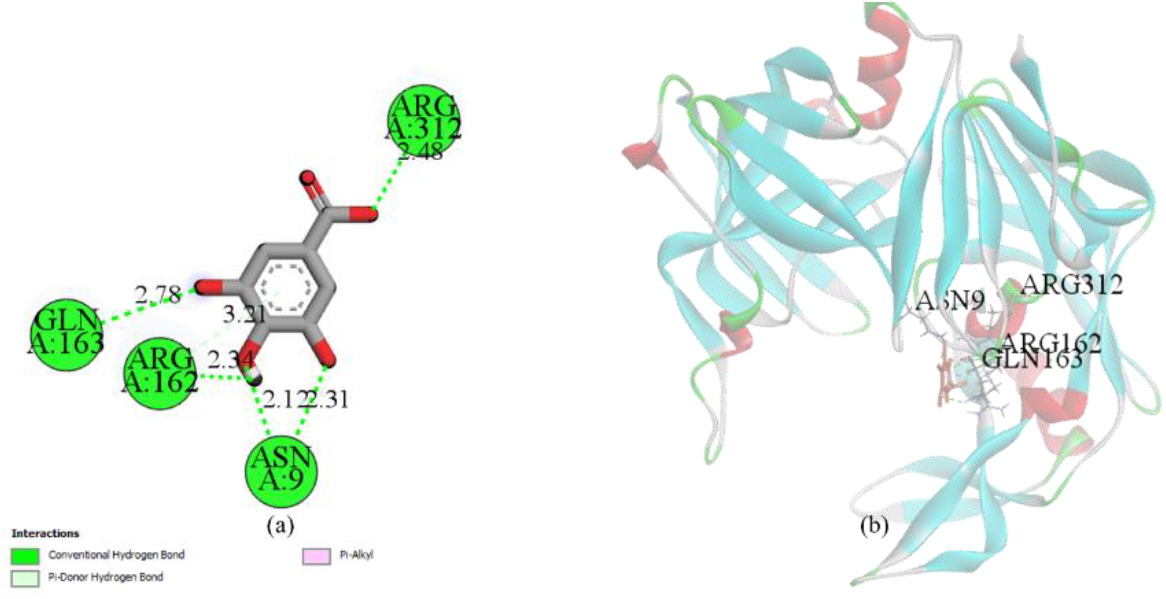
Visualization using BIOVIA Discovery Studio shows the interaction formed on the gallic acid-Sap 3 complex (a) 2D visualization shows bond distance, amino acid residues, and interaction type (b) 3D visualization shows semi-transparent complex

L-norepinephrine with the target receptor formed three electrostatic bonds, two of which have the same amino acid residue and ligand atom (Figure 6), which also formd hydrogen bonds, namely A:UNK1:H1 - A:ASP218:OD2 and A:UNK1:H3 - A: ASP218:OD2, while A:UNK1:N - A:ASP32:OD1 only formed electrostatic bonds. The distance created by the presence of hydrogen bonds ranges from 2.16417-5.09049 Å. The two catalytic residues formed hydrogen and electrostatic bonds with the ligands in this complex. The hydrophobic interactions formed in this complex are known to form in the amino acid residues VAL30 and ILE123, which had contact with the aromatic ring of the ligand. This ligand had contact with amino acid residues located at S1/S2/S3 and the catalytic residues.

**Figure 6.**
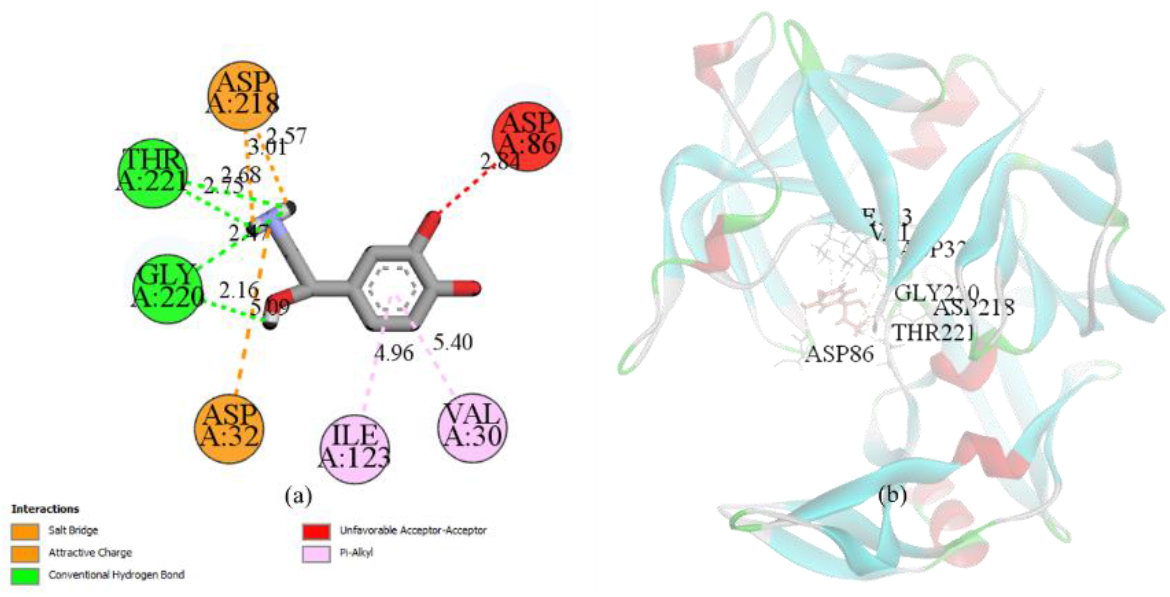
Visualization using BIOVIA Discovery Studio shows the interaction formed in the L-Norepinephrine-Sap 3 complex (a) 2D visualization shows bond distance, amino acid residues, and interaction type (b) 3D visualization shows semi-transparent complex

Mimosine also formed hydrogen bonds and electrostatic bonds on the identical amino acid residues and ligand atom (Figure 7), namely at A:UNK1:H1 - A:ASP218:OD2 and A:UNK1:H2 - A:ASP32:OD2. An electrostatic bond was also formed at A:UNK1:N - A:ASP32:OD1. Both receptor catalytic residues are formed in the two interactions. The distance created due to the presence of hydrogen bonds ranges from 1.99745 - 3.34731 Å. In this complex, the interactions formed at the VAL30, TYR84, and ILE123 amino acid residues had contact with the aromatic ring of the ligand. This ligand had contact with amino acid residues located at S1/S2/S3 and catalytic residues. It was reported that mimosine proved to be more efficient for controlling dermatophyte fungi in a previous study ^36^. Mimosine also has antimicrobial and antiviral activity ^36^.

**Figure 7.**
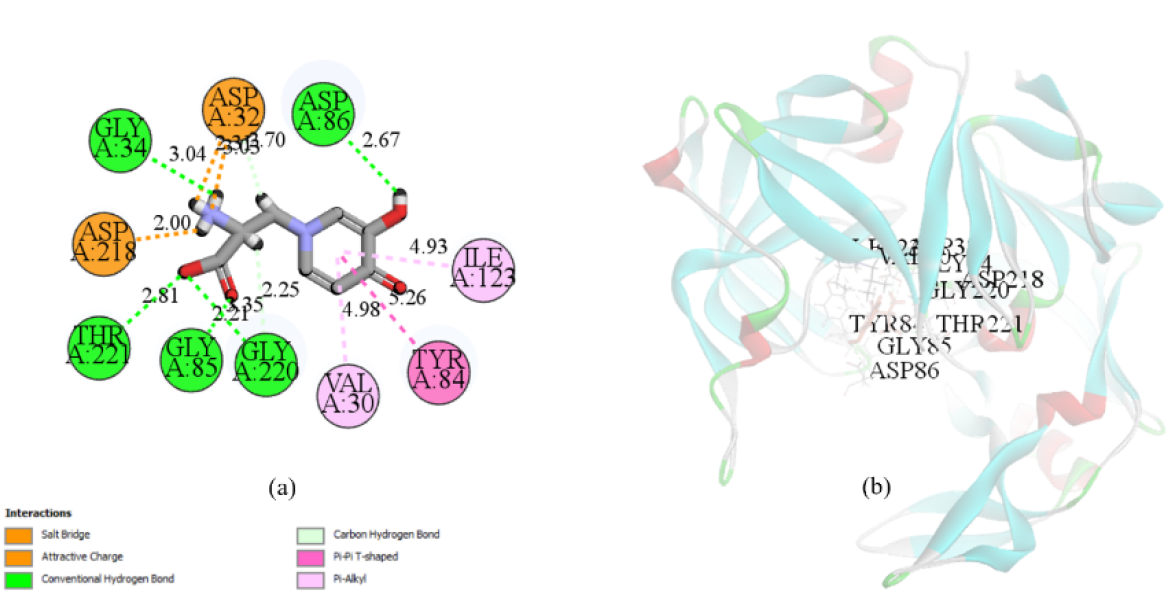
Visualization using BIOVIA Discovery Studio shows the interaction formed on the mimosine-Sap 3 complex (a) 2D visualization shows bond distance, amino acid residues, and interaction type (b) 3D visualization shows semi-transparent complex

Dominantly, L-ascorbic acid has hydrogen bond and one unfavorable bond with the target receptor (Figure 8). The distance generated by the presence of hydrogen bonds ranges from 2.12016 - 2.97822 Å. The O atom of the ligand binds with the amino acid residues GLY127, GLU193, and LEU194, while TYR128 forms the OH group. An unfavorable bond is formed between the amino acid residue ASN192 and the O atom of the ligand. The bond distance is 2.69808 Å with an unfavorable acceptor-acceptor type.

**Figure 8.**
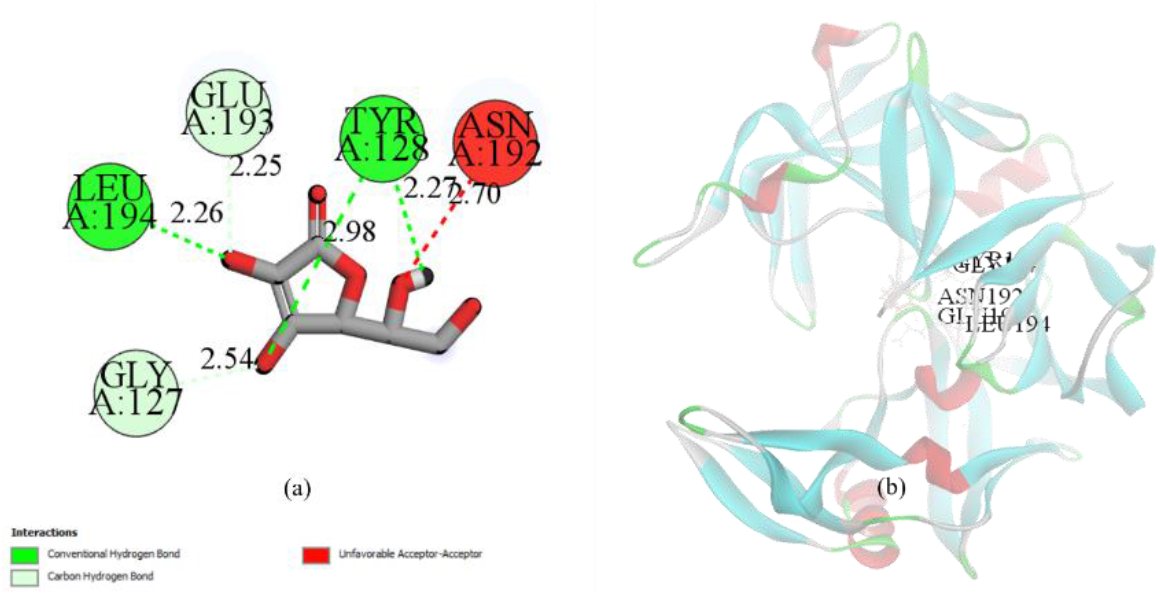
Visualization using BIOVIA Discovery Studio shows the interaction formed on the L-ascorbic acid-Sap 3 complex (a) 2D visualization shows bond distance, amino acid residues, and interaction type (b) 3D visualization shows semi-transparent complex

Linolenic acid has five hydrogen bonds and four hydrophobic interactions (Figure 9). The hydrogen bond distance formed ranges from 1.98702 - 3.19223 Å. The hydrogen bond in this ligand is also caused by the presence of from chemistry H-donor and to chemistry H-acceptor. THR222 and ILE223 bind with the ligand O atom.

**Figure 9.**
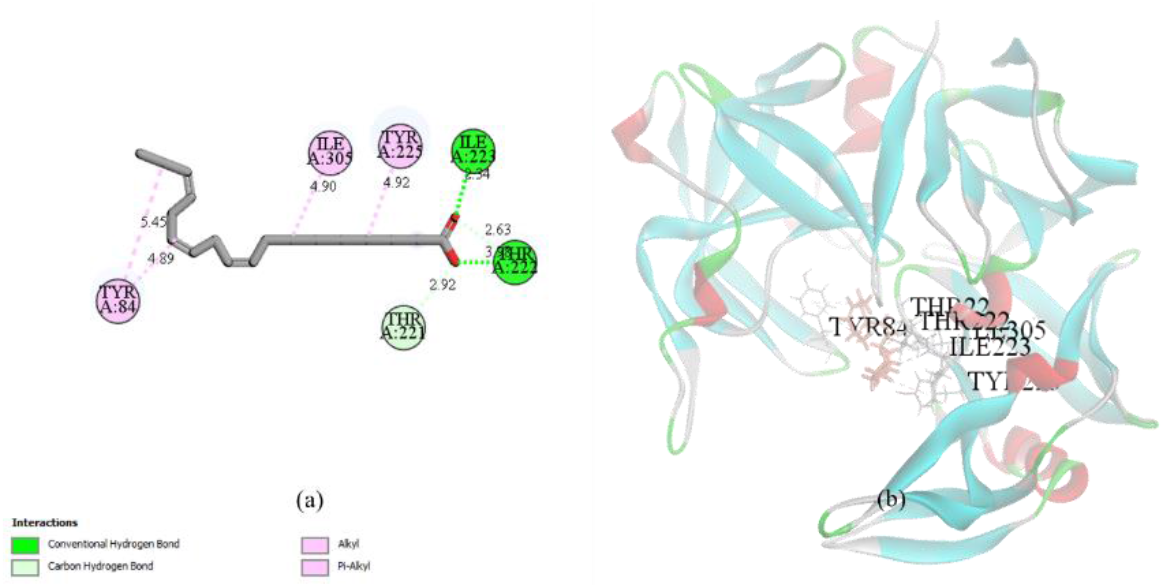
Visualization using BIOVIA Discovery Studio shows the interaction formed on the linolenic acid-Sap3 complex (a) 2D visualization shows bond distance, amino acid residues, and type of interaction (b) 3D visualization shows semi-transparent complex

The number of hydrogen bonds formed is known to determine binding strength in the receptor-ligand complexes. In addition, the efficiency of ligand binding to enzymes is also influenced by the binding energy ^30^. The phytochemicals present in medicinal plants are known to have the potential as antimicrobial and various other biological activities ^37^. All analysis data were summarized in Table 2.

We found in this study, the presence of unfavorable bonds were resulted from chemistry negative – to chemistry negative, from chemistry H-donor – to chemistry H-donor, and from chemistry H-acceptor – to chemistry H-acceptor. We hypothesized that the presence of such contacts in this study conferred instability in the receptor-ligand complex ^38^.

## 4. Conclusions

Based on the results of the present study, by utilizing the binding energy and interactions formed in the receptor-ligand complex and based on several other analytical parameters, we predict that the bioactive compounds of *Mimosa pudica* have a significantly good interaction with Sap 3, when compared with standard inhibitors. These data were completely structure-based prediction results, therefore further research are needed to strengthen and confirm the results of this study.

## Funding

This research received no external funding.

## Acknowledgments

This research has no acknowledgments.

## Conflicts of Interest

The authors declare no conflict of interest.

## Notes

### Competing Interest Statement

The authors have declared no competing interest.

